# MICA: Model-Informed Change-point Analysis

**DOI:** 10.64898/2026.03.16.712011

**Authors:** Mehdi Lotfi, Lars Kaderali

## Abstract

Change point detection is critical for identifying structural transitions in time series data. While most existing methods focus on changes in statistical properties of the data such as the mean or variance, many real-world systems are governed by dynamical models in which changes occur in model parameters. We introduce MICA, an algorithm that detects change points by minimizing the discrepancy between model simulations with a given dynamical model and observed data. The method integrates binary segmentation with a genetic algorithm to identify both the timing and nature of model parameter changes. MICA simultaneously estimates segment-specific and global parameters alongside change points, offering enhanced flexibility and interpretability. We demonstrate its utility on synthetic datasets and real-world scenarios, including COVID-19 epidemiological modeling, under policy interventions, and the analysis of generator cooling systems in wind turbines to monitor operational status. While illustrated using differential and difference equation models, MICA is model-agnostic and applicable to any simulatable system, making it broadly useful for applications requiring accurate tracking of structural dynamics.

## 1 Introduction

Time series models are foundational tools for analyzing dynamic systems across disciplines, capturing the sequential dependencies that drive temporal processes. From simple autoregressive models to complex dynamical systems described by differential equations, they are used extensively across scientific, engineering, and applied domains, ranging from physical and biological sciences to economics and environmental systems, to extract insights, forecast behavior, and inform decision-making.

A common challenge in time series analysis is the presence of structural transitions, or change points, where the underlying system behavior shifts due to external interventions or internal dynamics. Detecting such transitions from time series data, known as change point detection (CPD), is crucial for accurately characterizing system evolution, identifying anomalies, and improving forecasts. While some changes may coincide with known events (e.g., policy changes or hardware failures), others arise from latent processes, requiring automated and robust CPD methods.

The origins of CPD trace back to the foundational work of E.S. Page [1,2], who developed statistical techniques to detect distributional changes in sequential data. Since then, CPD has evolved into two broad classes: online methods, which detect changes in real time, and offline methods, which retrospectively segment entire datasets. Offline CPD has seen substantial growth, driven by applications in high-dimensional and nonlinear systems [3].

Recent advances have expanded CPD to specialized domains. In econometrics, methods based on cumulative sum statistics and score vectors capture regime shifts in volatility [4]. In neuroscience, modified binary segmentation and network-based approaches uncover changes in brain connectivity patterns [5, 6]. Other approaches leverage graphical models [8], kernel methods [9], Bayesian frameworks [7], and hidden Markov models [10] to capture changes in statistical properties such as mean, variance, or autocorrelation.

However, most CPD methods remain focused on statistical characteristics of the data, rather than the dynamics of the system generating it. This is a key limitation when analyzing time series governed by explicit mathematical models, such as differential equations. In such systems, structural changes often manifest as shifts in model parameters, not just in observable statistics. For example, during the COVID-19 pandemic, interventions like lockdowns and vaccination campaigns changed transmission dynamics, requiring updates to subset of the epidemiological model parameters.

This insight has motivated a growing interest in model-based CPD. Yet many existing methods impose restrictive assumptions, such as enforcing simultaneous changes in all parameters at each change point, or limiting the number of change points [11–13]. These simplifications can hinder both interpretability and accuracy in real-world applications.

To address these challenges, we introduce MICA (Model-based Change-point Analysis), a flexible algorithm designed to detect change points in time series data governed by an underlying, parametrized mathematical model. MICA frames CPD as a model selection problem, balancing goodness-of-fit with model complexity. It consists of two interacting components: a segmentation module, which proposes candidate change points using a modified binary segmentation strategy, and an optimization module, which estimates segment-specific and global parameters by minimizing the model-data discrepancy using a genetic algorithm.

A key innovation in MICA is its support for piecewise switching models, where both the locations of change points and the identities of changing parameters are inferred from the data, enabling only a subset of parameters to change across segments while others remain fixed. This enables fine-grained change point detection while maintaining continuity and model interpretability. The algorithm dynamically adjusts the parameter structure and leverages a cost function that penalizes over-segmentation, making it well-suited for both sparse and complex transitions.

We demonstrate MICA’s effectiveness on both synthetic and real-world datasets. First, we evaluate its performance under controlled conditions in a simulation study, varying noise levels, change point counts, data lengths and so on. Second, we apply MICA to two case studies: (1) modeling the spread of COVID-19 in Germany using an ODE-based compartmental model, and (2) detecting operational status changes in wind turbine cooling systems using difference equations and Supervisory Control and Data Acquisition (SCADA) data. In both applications, MICA identifies interpretable change points that correspond to known interventions or system events, while revealing subtle transitions missed by traditional methods.

The remainder of this paper is organized as follows. Section 2 introduces the mathematical framework of MICA, including the piecewise model structure and the underlying optimization strategy. Section 3 presents results from benchmark experiments and real-world applications. We conclude with a discussion of the method’s strengths, limitations, and potential future directions.

## 2 Methodology

MICA is a model-based change point detection framework designed for time series with underlying mathematical models. It identifies change points by minimizing the discrepancy between model simulations and observed data, while simultaneously estimating global (non-segment-specific) and local (segment-specific) model parameters. MICA supports scenarios in which only a subset of parameters changes across time, making it suitable for complex systems with large numbers of parameters, only some of which are allowed to change over time.

Although this section presents MICA in the context of ordinary differential equation (ODE) models, the framework is model-agnostic and applicable to any time series system defined by explicit mathematical formulations. This includes discrete-time systems, algebraic or hybrid models, stochastic processes, and other frameworks. The only requirement is that the model can be simulated over individual time segments, has parameters governing its behavior, and produces output that can be compared to observed data.

### 2.1 Problem Formulation

#### Notations

- Let 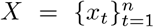 represent a time series of length *n*. A segment 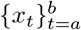 is a contiguous subset satisfying 1≤ *a < b*≤ *n*. We denote the full dataset as *x*_1…*n*_ and subsegments as *x*_*a*…*b*_.
- A set *CPs* ={ *t*_1_, *t*_2_, …, *t*_*k*_} ⊂ {1, 2, …, *n*},with *t*_1_ *< t*_2_ *<*… *< t*_*k*_, identifies the indices of change points, dividing the dataset into *k* + 1 segments. These points correspond to changes in some of the parameters of the governing model.

MICA formulates change point detection as a model selection problem in the context of time series governed by parametric models. In many real-world systems, certain parameters governing system behavior may remain stable over time, while others can experience abrupt changes at unknown points due to internal dynamics or external interventions. Identifying these change points and accurately estimating model parameters across segments is critical for understanding system dynamics.

We use ordinary differential equations (ODEs) as a reference modeling framework. A classical ODE system is defined as:

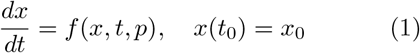

where *x* denotes the vector of state variables, *t* is time, and *p* is a set of fixed model parameters. This standard formulation assumes the dynamics of the system are invariant throughout the observed period.

To incorporate structural changes, we extend this to a piecewise formulation, Piecewise Switching ODEs (PSODEs):

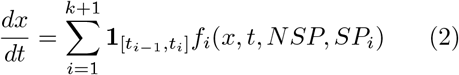

Here, *k* is the number of change points dividing the time horizon into *k* + 1 segments. Each segment [*t*_*i−*1_, *t*_*i*_) is governed by a potentially distinct parameterization of the model. *NSP* denotes global (non-segment-specific) parameters shared across all segments, while *SP*_*i*_ are segment-specific parameters local to the *i*-th segment.

The initial conditions for the PSODE are defined recursively. The system starts from the given initial state *x*(*t*_0_) = *x*_0_ for the first segment, and the state at the end of each segment serves as the initial condition for the next:

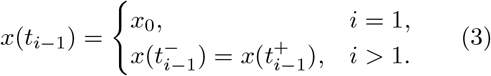

This ensures continuity of the system across change points.

Also, 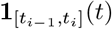 is the indicator function defined as:

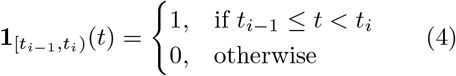

Let Θ = *NSP* ∪ *SP* denote the full set of model parameters, where *NSP* represents non-segment-specific parameters shared across all segments, and *SP* = {*SP*_1_, …, *SP*_*k*+1_} denotes the segment-specific parameters associated with each segment.

Formally, the objective is to estimate the set of change points *CP* = {*t*_1_, …, *t*_*k*_}together with the model parameters Θ by minimizing the following penalized cost function:

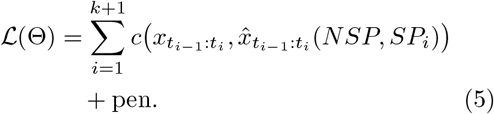

Here, 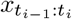 denotes the observed data on segment *i*, and 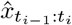 (*NSP, SP*_*i*_) is the corre-sponding model output obtained by simulating the piecewise dynamical system on that segment using the global parameters *NSP* and the local segment-specific parameters *SP*_*i*_. The function *c*(·,·) denotes a segment-wise discrepancy measure (e.g., squared error or RMSE), and pen is a penalty term that controls over-segmentation (e.g., a BIC-based penalty).

Minimizing ℒjointly determines the number and locations of change points, the global parameters shared across all segments, and the segment-specific parameters, while balancing goodness of fit against model complexity.

In this study, we employed a BIC-based penalty, which provides a principled trade-off between model fit and segmentation complexity and was applied consistently across all analyses.

### 2.2 MICA Architecture

MICA performs change point detection by iteratively refining a piecewise model that best explains the observed time series. It begins with the entire dataset modeled as a single segment and sequentially introduces candidate change points. At each step, data and model are scanned using a sliding window approach to identify positions where splitting the time series leads to a meaningful improvement in model fit. For each proposed split, parameters are estimated across all segments, and the global model error is evaluated. New change points are accepted only if they reduce the overall discrepancy between the model simulations and observed data by more than the penalty value, effectively balancing model accuracy and complexity. As new segments are introduced, the model’s parameter structure dynamically adapts, ensuring consistency across the full time course.

The algorithm proceeds through iterative scans in both forward and backward directions to avoid missing any relevant structural changes. This process is stack-wise: after each newly detected change point, the resulting segments are added to a stack of segments awaiting evaluation. The forward check begins by evaluating the last segment for a potential change point. If no change point is detected, the segment is removed from consideration, and the algorithm moves to the second-to-last segment, this is the backward check phase. This backward evaluation continues until either a change point is found or all segments have been examined.

If a new change point is detected in the backward pass, the algorithm performs another forward and backward scan from the newly created segments to ensure that no significant change points are overlooked.

The final result is a segmented model in which each interval reflects a distinct dynamic regime, characterized by its own parameter subset. This process is illustrated in the MICA flowchart shown in Figure 2, where red arrows denote the initial iteration without any change points, and black arrows indicate subsequent refinement steps.

**Fig. 1:**
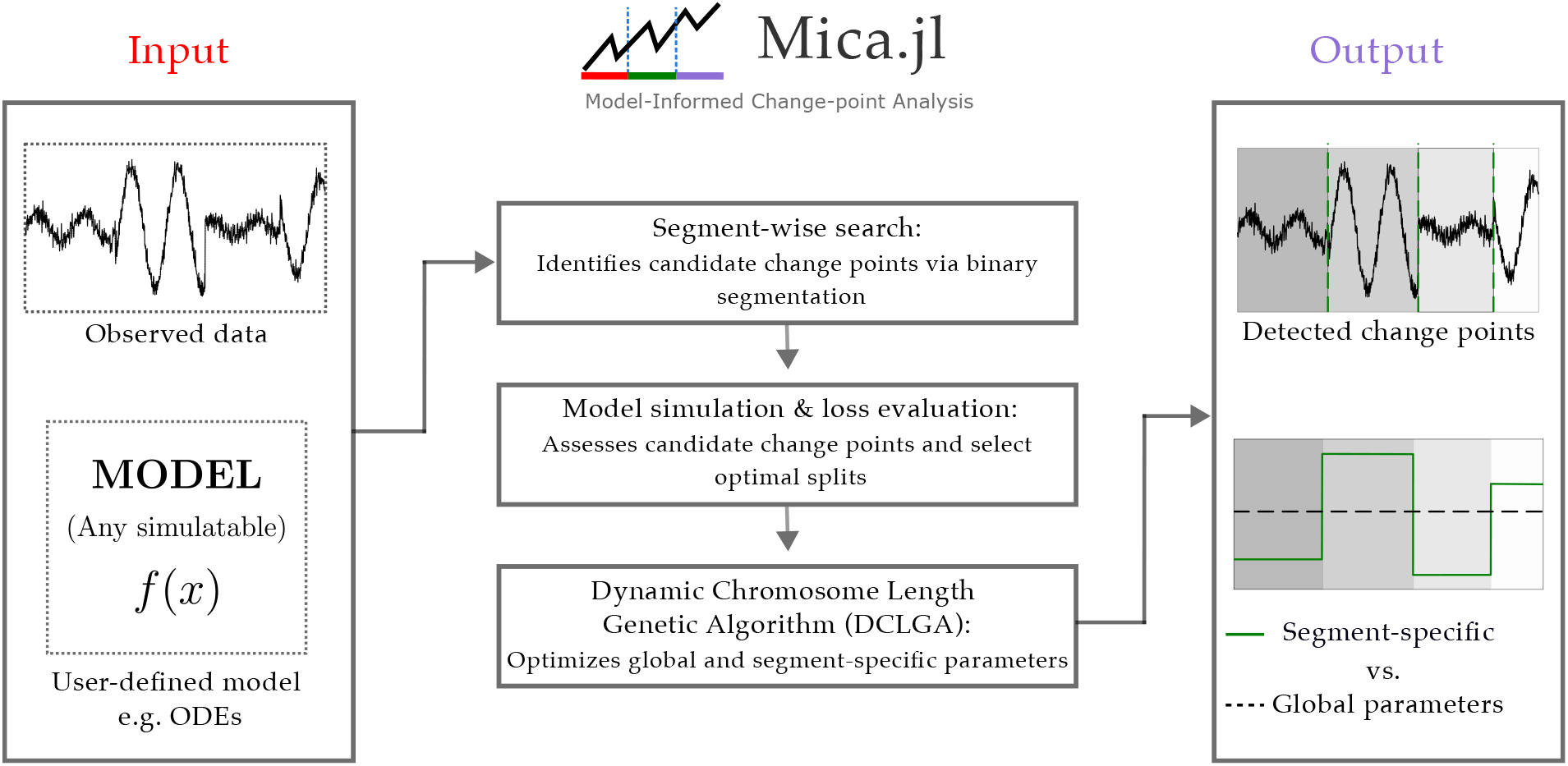
Graphical Abstract

**Fig. 2:**
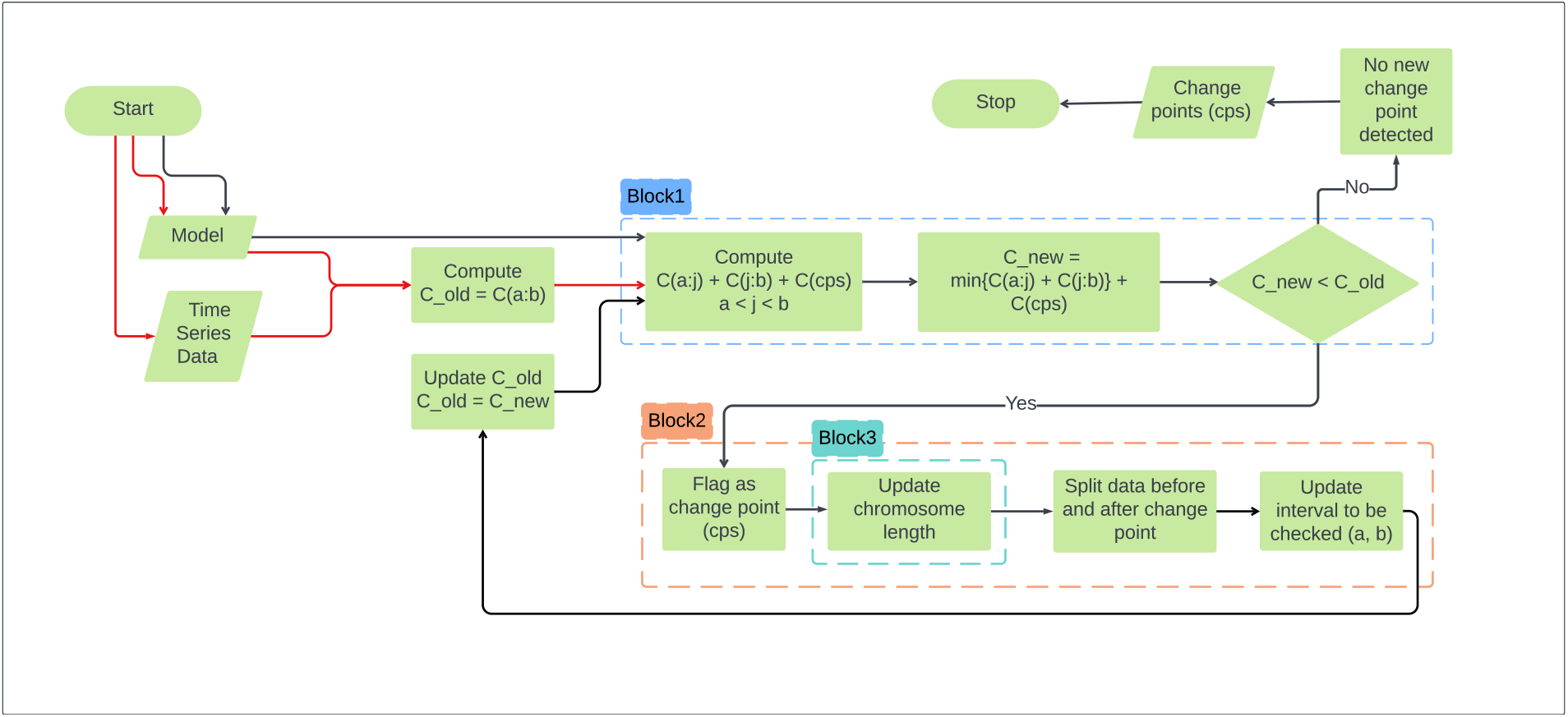
MICA algorithm overview. The algorithm starts with an initial model and dataset, and iteratively proposes and validates change points. The framework integrates three key modules: segmentation (Block 2), optimization (Block 1), and chromosome updates (Block 3). Red arrows indicate the initialization step without change points; black arrows show subsequent iterations.

As described in Algorithm 1, the MICA framework uses a genetic algorithm to search and jointly optimize global and segment-specific parameters, while iteratively identifying change points that minimize the penalized model-data discrepancy. The pseudocode summarizes this iterative process, including candidate evaluation, model simulation, and dynamic chromosome updates, which together form the core logic behind the segmentation, optimization, and interaction modules.

#### Algorithm 1

MICA algorithm for detecting change points in dynamical systems by minimizing simulation-based loss using genetic optimization.

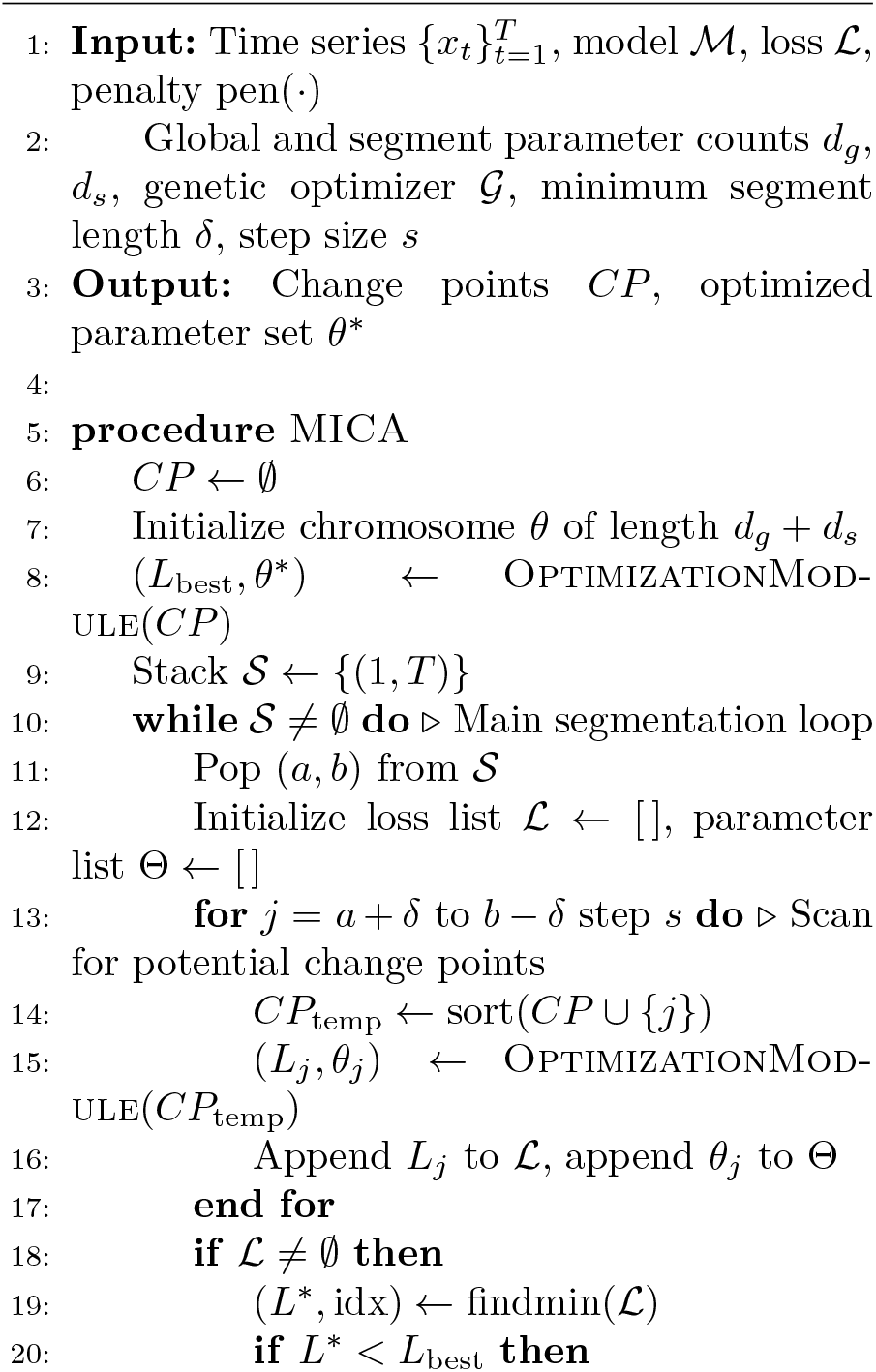
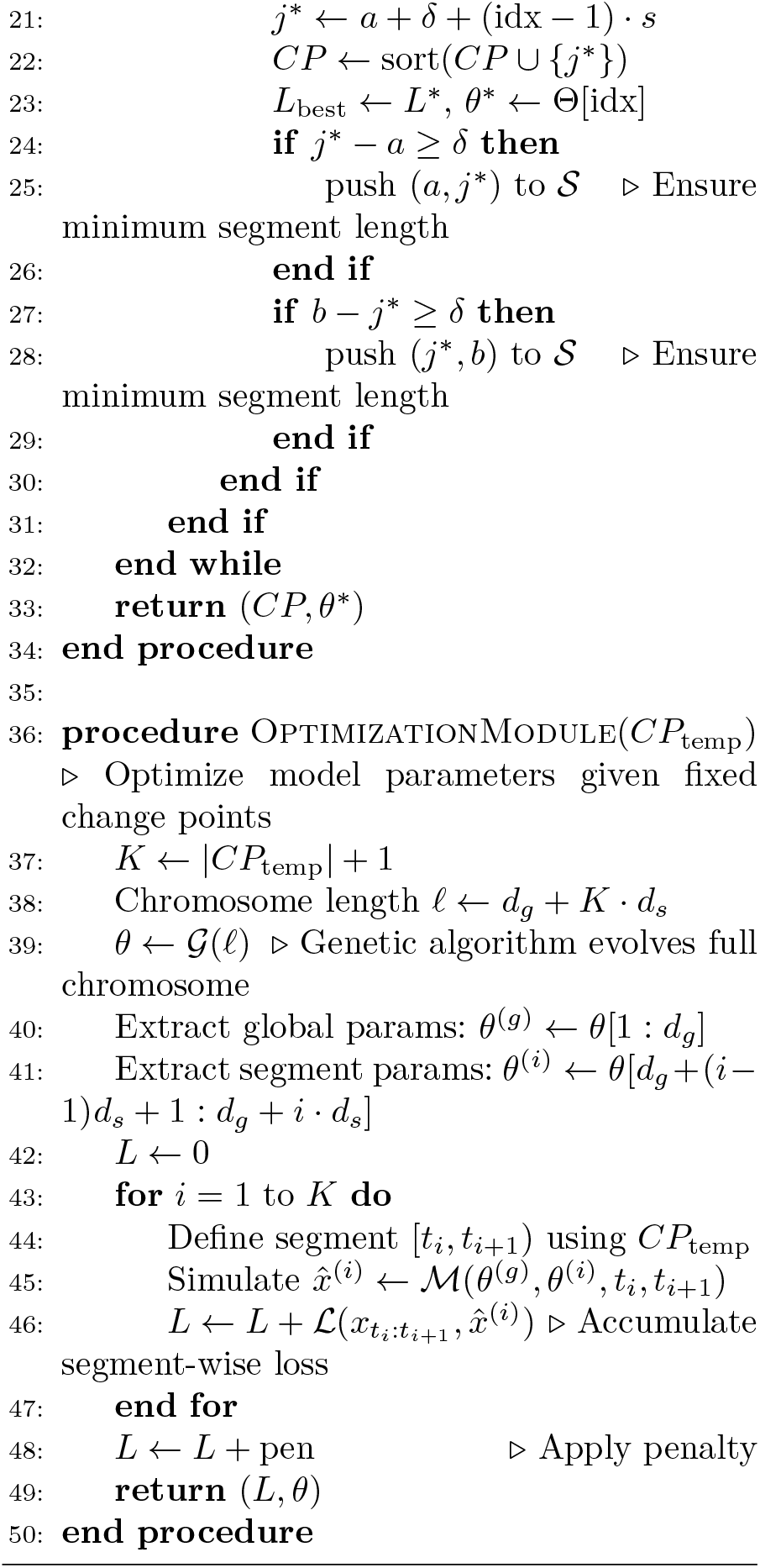

#### 2.2.1 Segmentation Module

In this section, we present the Segmentation Module, a core component of the proposed change point detection framework. Inspired by classical binary segmentation, this module iteratively partitions the time series into smaller segments and identifies candidate change points, which are then passed to the Optimization Module for validation (described in the next subsection).

To tailor binary segmentation for our model-based approach, we modified the traditional cost function, originally based on statistical characteristics, to incorporate outputs from ODE simulations alongside observed data. This adaptation enables the algorithm to evaluate how well each segment captures the underlying system dynamics, rather than relying solely on statistical patterns.

By incorporating simulation-based model behavior into the segmentation process, MICA ensures that detected change points correspond to meaningful shifts in the system’s dynamics.

Figure 3 illustrates the detailed workings of the segmentation process. The procedure begins by analyzing the entire dataset to identify the first potential change point. A counter with a sliding window, denoted by *j*, identifies potential change points and the associated segments, which are then passed to the Optimization Module. The graphical representation of the counter *j*, which shows the index of data points at a given time point, is depicted in the upper part of Figure 3. The process is initialized with a minimum segment length of 10 data points. During each iteration, the counter increments by one data point, progressing according to the expression *j* = *a* + 10 : 1 : *b*− 10 where *a* and *b* represent the start and end indices of the current segment. Initially, *a* is set to 0 and *b* to n, the total length of the given dataset. Enforcing this minimum segment length constraint ensures that each segment contains sufficient data to enable accurate parameter estimation during the optimization stage. This condition guarantees that each partition contains at least 10 time points, which is essential for accurate parameter estimation in the Optimization Module. For example, fitting ODEs with an insufficient number of data points would pose significant challenges.

**Fig. 3:**
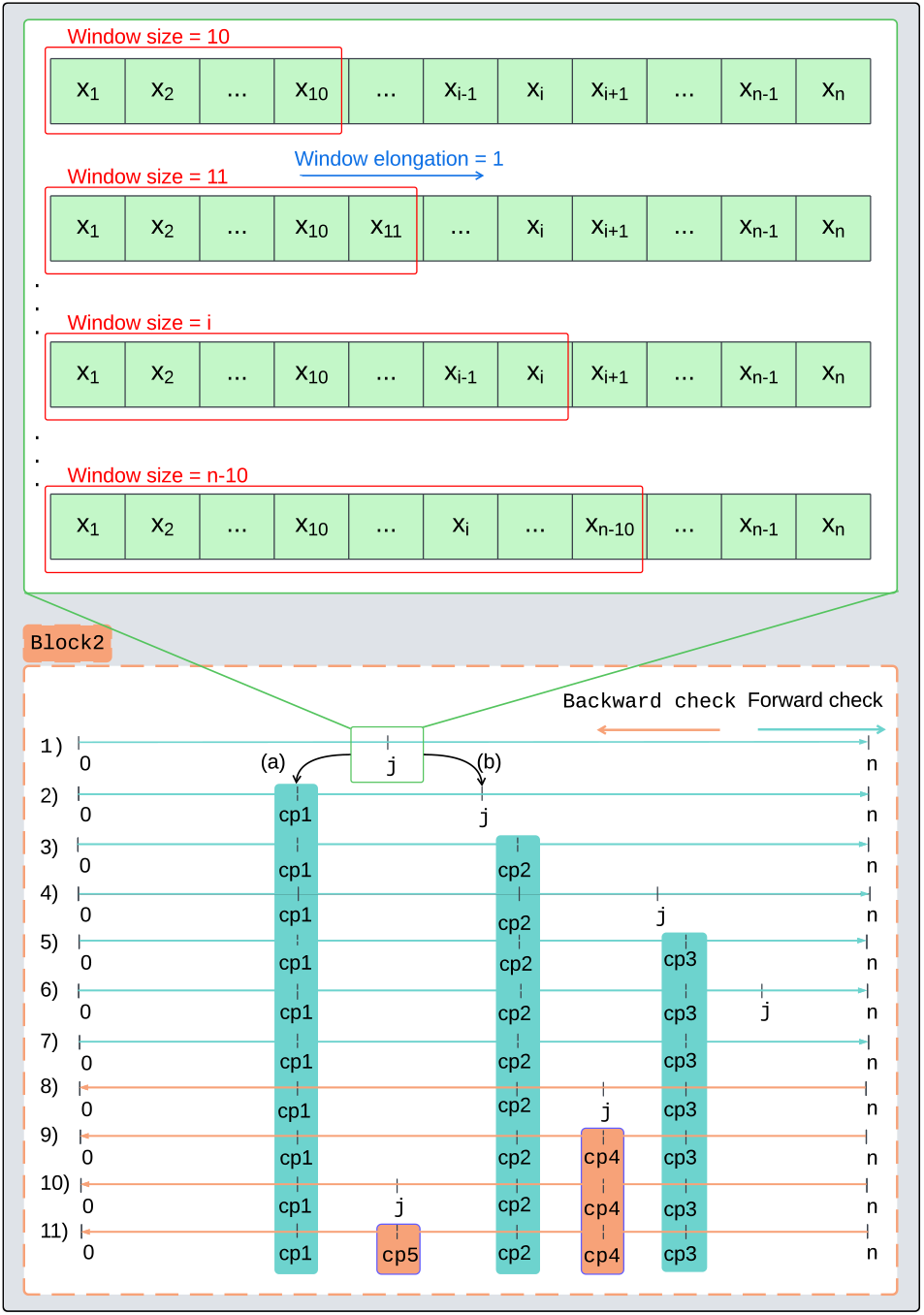
Segmentation Module: Illustration of the forward-backward traversal logic using a sliding window. The algorithm incrementally updates the counter *j* to identify optimal segment splits while enforcing minimum segment length.

During the first iteration (lower part of Figure 3), the Optimization Module performs two tasks: (a) it identifies the optimal position for *j* that minimizes the total loss between the observed dataset and the model, designating this position as *cp*_1_; and (b) it updates the interval for the counter *j*. Once the first change point is identified, the algorithm proceeds with subsequent iterations by fixing the initial change point and focusing the counter on the last segment created by this change point to search for the next potential change point. This process is referred to as the forward check.

In subsequent iterations (2-7), after identifying the first change point, the algorithm continues using the forward check (indicated by light blue arrows in Figure 3) to Explore additional potential change points until it reaches the end of the dataset. In each iteration, the algorithm focuses on the most recent segment created by previously detected change points, leaving the remaining segments for later analysis. Any segments not immediately checked are deferred for subsequent analysis. Once the forward check is completed, the algorithm initiates a backward check (iterations 8-11). During the backward check, the algorithm revisits previously skipped segments to identify any additional change points that may have been missed during the forward check. For example, in iteration 7, since no change point was identified between *cp*_3_ and the end of the dataset, the algorithm places the counter between *cp*_2_ and *cp*_3_ (iteration 8), resulting in the identification of *cp*_4_ (iteration 9). In iteration 10, it is assumed that there is no change point between *cp*_4_ and *cp*_3_, and also between *cp*_2_ and *cp*_4_; otherwise, a forward check would have been required for the former case and a backward check for the latter. The detailed computations, particularly the cost calculations for a given interval, are elaborated in the Optimization Module subsection.

In summary, the Segmentation Module in MICA generates potential change point locations, which are then passed to the subsequent Optimization Module for further refinement and validation.

#### 2.2.2 Interaction Module

The Interaction Module (Figure 4) serves as the central component that coordinates the Segmentation and Optimization Modules to ensure coherent and accurate change point detection. Once potential change points are identified by the Binary Segmentation (BS) module, the dataset is divided into segments accordingly. For each segment, the system of ordinary differential equations (ODEs) is solved over the segment’s time span, producing simulated outputs that represent the system’s expected behavior during that interval. To maintain dynamic consistency across the time series, the final state of each segment’s solution is used as the initial condition for the next segment.

**Fig. 4:**
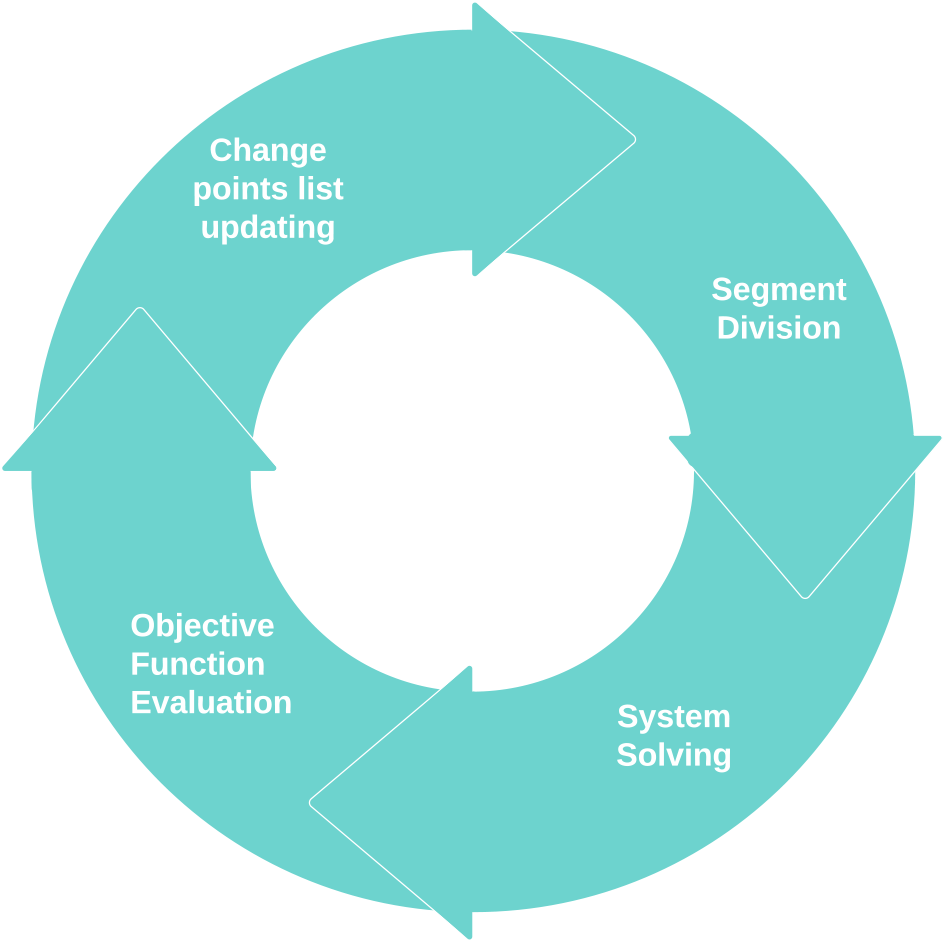
Interaction Module: This module coordinates the other two modules, namely the segmentation and optimization modules, through four steps. A detailed explanation of each step is provided in the text.

Subsequently, an objective function (defined in Equation 5) is evaluated by comparing the simulated outputs to the observed data within each segment. This function quantifies the discrepancy between the model and the data and also incorporates a regularization mechanism that discourages overly complex models, ensuring that the identified change points are both accurate and interpretable.

By combining the initial segmentation from the BS module with a rigorous ODE-driven optimization framework, this module guarantees that the final set of change points is both robust and meaningful, capturing genuine shifts in the system’s behavior. The two-step process leverages the strengths of both modules, resulting in a comprehensive and data-driven approach to ODE-based change point detection.

#### 2.2.3 Optimization Module

The Optimization Module takes as input the potential change points identified by the Segmentation Module and uses them to partition the dataset into corresponding segments. For each segment, the module solves the specified system of ordinary differential equations (ODEs) over the appropriate time interval, using the segment’s observed data and temporal boundaries. The simulated outputs from the ODE system are then incorporated into an objective function that quantifies the fit between the model and the empirical data within each segment, ensuring a detailed and accurate assessment of model performance.

Given the interdependence of segments introduced by change points, the objective function is computed cumulatively. At each iteration of change point detection, the total cost is calculated as the sum of individual costs from all segments, both those established in previous iterations and those defined by new candidate change points. This cumulative approach is essential for capturing the global impact of adding or refining change points, allowing the optimization process to evaluate whether the introduction of a new change point improves the model’s overall explanatory power. By considering the entire dataset in this aggregated manner, the algorithm effectively identifies change points that correspond to genuine shifts in the underlying system dynamics.

Figure 5 illustrates the Optimization Module, which is responsible for detecting change points within a given segment during each iteration of the broader change point detection process. Our method for identifying change points in time series data relies on an iterative segmentation strategy that locates time points where the system dynamics undergo significant changes. To efficiently manage which segments are yet to be analyzed, we maintain a dynamic list that is updated throughout the procedure. The implementation of this is inspired by Julia’s built-in `push!` and `pop!` commands, which are commonly used for manipulating elements in arrays or stacks. Specifically, `push!` appends a new segment to the list, while `pop!` removes and retrieves the most recently added segment. This stack-based mechanism allows the algorithm to systematically explore and process each segment, ensuring thorough coverage of the dataset in the search for change points.

**Fig. 5:**
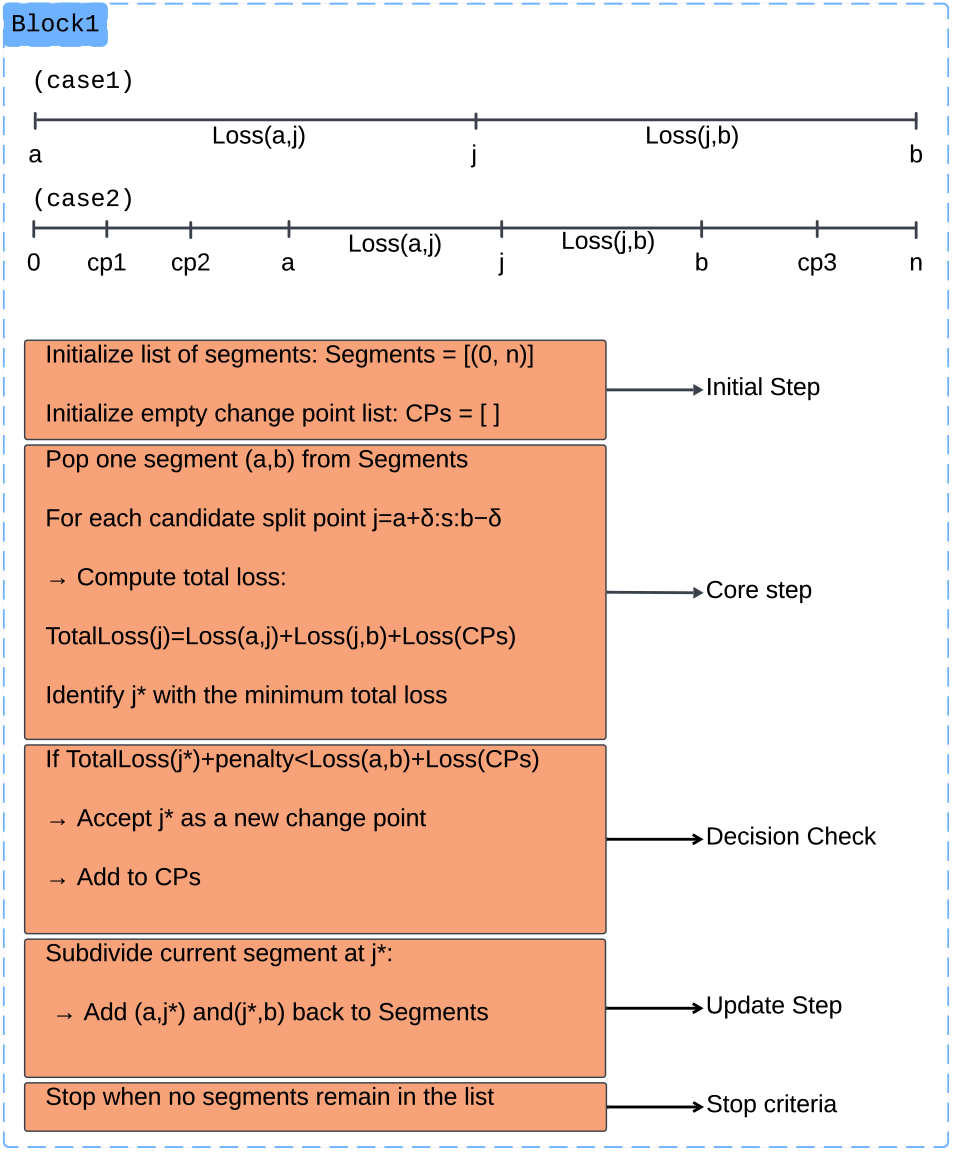
The change point detection process within a segment during an iteration of the overall procedure. Initially, the start and end of the dataset are considered, with a counter representing the first potential change point. As the process progresses, detected change points define new segments. Costs associated with different values are computed and stored. The minimum cost and its index are identified and, if inserting this change point reduces the overall cost, it is designated as a new change point. The segment is then split, and operations are applied to the new segments. The algorithm stops when there are no more segments to investigate.

In the **initial step**, the algorithm begins by preparing the core data structures required for change point detection. A variable (`Segments`) is initialized to store the segments to be analyzed, starting with a single segment that spans the entire dataset, from index 0 to *n*, where *n* is the dataset length. This initial segment is added to the list of intervals scheduled for processing. Simultaneously, another variable (`CPs`) is prepared to keep track of the change points identified during the process. This list is initially empty and grows as the algorithm iteratively uncovers new points of change. In the **core step**, the algorithm retrieves the most recently added segment, denoted (*a, b*), from the list of segments awaiting analysis. This represents the current interval under investigation. In the first iteration (Case 1 in Figure 5), *a* and *b* correspond to the start and end of the dataset. As change points are progressively discovered, subsequent iterations operate on smaller intervals defined by previously identified points. As explained in Segmentation Module section, a counter *j* is introduced within this segment to evaluate potential positions where a change might have occurred. To maintain the reliability of parameter estimation, the counter is constrained to examine positions that yield at least *δ* data points in each resulting subsegment. Thus, *j* takes values from *a* + *δ* to *b* − *δ*, where *δ* = 10 in our implementation and the step size is set to *s* = 1. These values can be adjusted depending on the specific characteristics of the problem. For each candidate *j*, the algorithm computes the total cost of dividing the segment at that point. This cost includes the sum of the losses for the proposed subsegments (*a, j*) and (*j, b*), along with the cumulative cost associated with segments formed by previously accepted change points. These values are stored in a temporary list of total costs. Case 2 in Figure 5 illustrates this process in action: after detecting several change points (e.g., *cp*_1_ through *cp*_3_), the algorithm examines a smaller subinterval, sweeping the counter *j* to test potential split locations. The algorithm then selects the value of *j* that results in the minimum cost (*j*^*\**^). If the newly computed minimum cost is lower than that of the previous configuration, the associated *j* is confirmed as a new change point. This value is appended to the growing list of detected change points. In the **update step**, once a new change point *cp* is validated, the current segment (*a, b*) is divided into two subsegments: (*a, cp*) and (*cp, b*). These new intervals are then queued for further evaluation, allowing the algorithm to recursively analyze increasingly refined sections of the dataset. The process continues iteratively: at each step, segments are retrieved from the queue, analyzed, and if warranted, further subdivided based on newly discovered change points. The algorithm halts when there are no more segments left to examine, specifically when the list of segments is empty, signaling that all meaningful structural shifts in the data have been detected.

In summary, the algorithm dynamically manages the segmentation process through the use of segment insertion and removal operations, iteratively refining the detection of change points by focusing on progressively smaller intervals until no further division is warranted.

#### 2.2.4 Chromosome Encoding and Update

In the Optimization Module, a Dynamic Chromosome Length Genetic Algorithm (DCLGA) is employed to minimize an objective function that quantifies the discrepancy between the model simulations and the observed time series data. As visualized in Figure 6 the algorithm operates on a fixed-length chromosome that encodes both global (segment-independent) parameters, denoted as *a*_1_, *a*_2_, …, *a*_*m*_, and segment-specific parameters, denoted as *b*_1_, *b*_2_, …, *b*_*n*_, where *m* is the number of global parameters and *n* is the number of parameters specific to each segment.

**Fig. 6:**
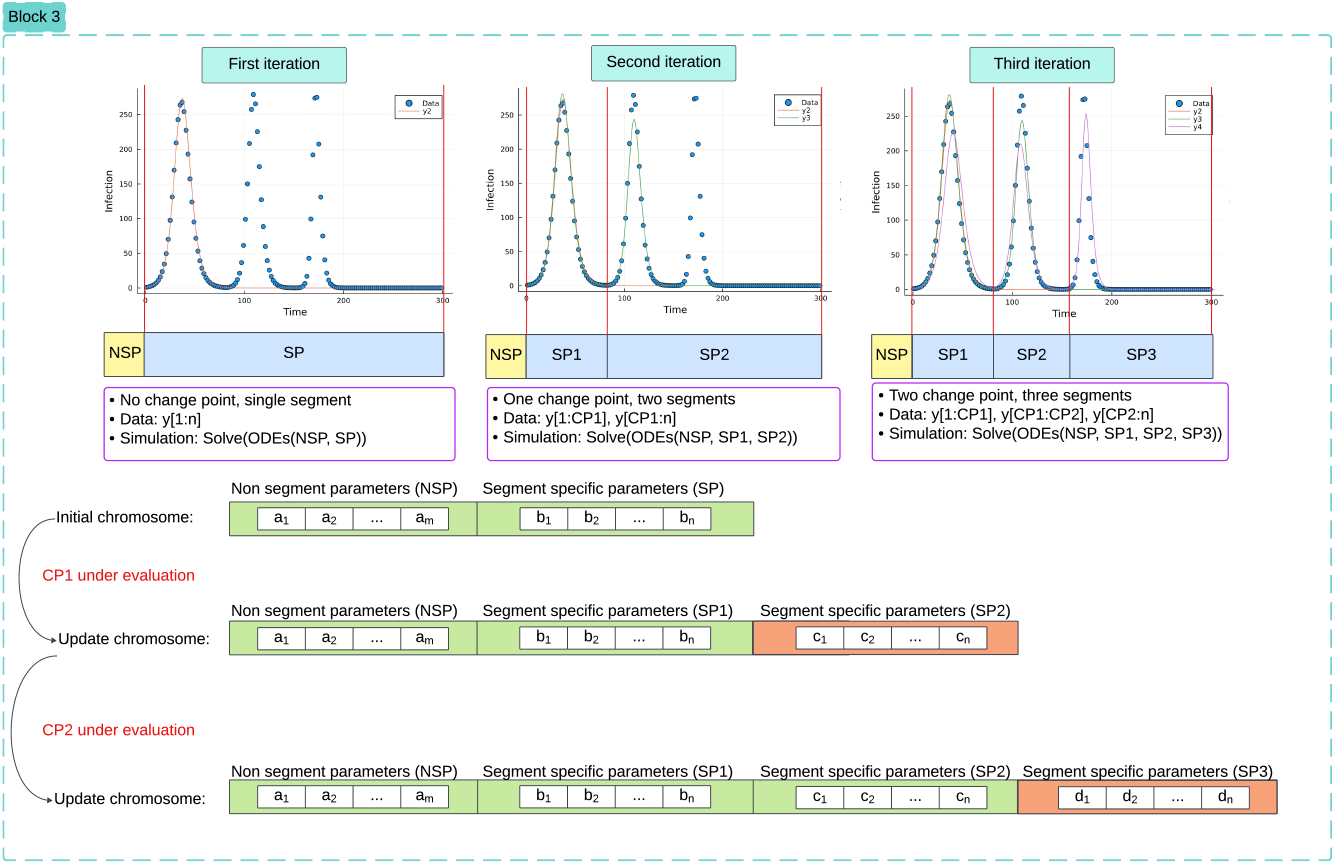
Dynamic Chromosome Length Genetic Algorithm (DCLGA). The top row illustrates three iterations of the change point detection process applied to a toy dataset generated from a system of ODEs. Each iteration highlights the number of detected change points, the corresponding segmentation of the data, and the solution of the model, which incorporates both segment-specific and segment-independent parameters. The bottom row depicts the mechanism for updating the chromosome structure within the genetic algorithm as new change points are introduced.

Initially, in the absence of any change points, the entire time series is treated as a single segment. The genetic algorithm then optimizes the parameter set corresponding to this configuration. A counter *j* is introduced to identify the first change point (as detailed in the Segmentation and Optimization Modules). Once a change point is detected, the time series is split into two segments. The chromosome is accordingly extended to include a second set of segment-specific parameters, denoted as *c*_1_, *c*_2_, …, *c*_*n*_, while the global parameters *a*_1_, …, *a*_*m*_ remain unchanged and are still positioned at the beginning of the chromosome.

The change point detection process continues iteratively. After the first change point is established, the algorithm proceeds to search for a second change point using a chromosome of the same current length. When the second change point is identified, the chromosome is again expanded to accommodate a third segment with its own parameters *d*_1_, *d*_2_, …, *d*_*n*_, maintaining the same structure.

At each iteration, the chromosome comprises three main components: (1) the global parameters shared across all segments, (2) the optimized parameters for previously identified segments, and (3) the initial guesses for the parameters of the newly proposed segment. This evolving structure allows the genetic algorithm to efficiently adapt the model to increasingly complex segmentations while preserving continuity and interpretability of parameter roles.

### 2.3 Implementation

We provide an open-source Julia package for MICA, supporting modular model definitions and user-defined loss functions. The initial release includes built-in support for ODE-based models and difference equations. The package is extensible to custom mathematical formulations and includes tools for model simulation and parameter configuration.

Comprehensive documentation and tutorials are available on the MICA project website (https://changepointdetection.com), the GitHub repository (https://github.com/Mehdilotfi7/MICA), and the institute’s official software page (https://wordpress.kaderali.org/software/).

## 3 Results

In this section, we evaluate MICA on synthetic datasets and in real-world applications. We begin by evaluating the algorithm on synthetic datasets generated from ordinary differential equations (ODEs) under various controlled conditions in a simulation study, as described in the subsequent subsection. These synthetic scenarios enable a rigorous assessment of the algorithm’s performance, specifically its accuracy in detecting change points, robustness to noise and parameter variations and run time, as true change points and parameters are known.

All simulations and benchmarking experiments were conducted on a Lenovo ThinkStation P2 Tower equipped with an Intel^®^ Core™ i9-14900K processor (32 threads), 64 GiB RAM, and an NVIDIA GeForce RTX™ 4060 GPU. The system ran Ubuntu 24.04.3 LTS (64-bit) with Linux kernel 6.14.0-29-generic and GNOME 46 (X11).

Following the synthetic evaluation, we demonstrate the algorithm’s applicability to real-world datasets in two distinct domains. The first application involves change point detection in epidemiological model simulations, aimed at analyzing the effects of public health interventions on the spread of COVID-19 in Germany. This case study addresses two key objectives: first, to identify the timing of change points corresponding to events such as policy interventions and behavioral shifts; and second, to quantify the resulting impact on model parameters governing disease transmission. By pinpointing when these changes occurred and how they affected system dynamics, we gain insights into the effectiveness of different interventions. The second application focuses on operational data from wind turbines and a generator cooling system, showcasing the algorithm’s utility in the renewable energy sector. In this context, the algorithm is employed to detect operational changes, thereby supporting condition monitoring and predictive maintenance strategies.

### 3.1 MICA in Synthetic Datasets

To assess the performance of MICA, we first apply it to synthetic time series data generated using the SIR (Susceptible–Infectious–Recovered) model, a classical compartmental framework in epidemiology that describes the flow of individuals through three states: susceptible to infection, actively infectious, and recovered (and thus immune). This model captures the core dynamics of epidemic spread and is a widely used approach to model infectious diseases. In our analysis, only the time series of the infectious compartment is used in the objective function for parameter estimation, reflecting a common scenario where limited observational data is available. Our aim here is to test the MICA’s ability to detect structural changes in model parameters, specifically, changes in the transmission rate (*β*) while holding the recovery rate (*γ*) constant.

This setup reflects realistic scenarios in which public health interventions, behavioral changes, or environmental factors can alter transmission dynamics, while recovery rates tend to remain stable over time. The temporal evolution of the disease is governed by the following system of coupled ordinary differential equations (ODEs):

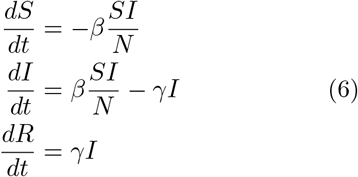

where *S*(*t*) denotes the number of susceptible individuals at time *t, I*(*t*) denotes the number of infectious individuals at time *t, R*(*t*) denotes the number of recovered individuals at time *t, β* is the transmission rate of the disease, which is assumed to vary by segment, *γ* is the recovery rate, considered constant across segments, and *N* represents the total population size, given by *N* = *S*(*t*) + *I*(*t*) + *R*(*t*).

#### 3.1.1 Experimental Design and Criteria

To evaluate the effectiveness of MICA, we generate synthetic (toy) datasets based on a comprehensive set of criteria, as detailed in Table 1. The algorithm was then applied to each generated dataset. The evaluation criteria include variations in the number of change points, dataset lengths,types and levels of noise, and penalty values. For each specific combination of these parameters, a corresponding dataset is generated to test the robustness and accuracy of the algorithm.

**Table 1:** Experimental design and evaluation criteria for change point detection. This table summarizes the experimental design used to assess the robustness of the proposed change point detection method across a range of conditions, including the number and locations of change points, dataset lengths, noise types and levels, and penalty values. Synthetic datasets are generated according to these criteria and used to systematically evaluate detection accuracy and computational performance.

| Evaluation Criterion | Values |
| --- | --- |
| Number of Change Points | {1, 2, 3} |
| Data Lengths | {70, 130, 160, 200, 250} |
| Noise Types | {"Gaussian", "Uniform"} |
| Noise Levels | {1.0, 10.0, 20.0} |
| Penalty Coefficient ($\kappa$) | {0.0, 1.0, 10.0, 20.0, 30.0, 100.0} |
| $\beta$ Values | {0.00009, 0.00014, 0.00025, 0.0005} |
| $\gamma$ Value | 0.7 |
| Change Points Positions | {50.0, 100.0, 150.0} |

The toy dataset is generated using the SIR model equations described in Equations 6, incorporating two change points located at time indices 50 and 100, which divide the dataset into three distinct segments. Segment-specific parameter values for the transmission rate *β* (while keeping the recovery rate *γ* fixed) are assigned according to the configuration specified in Table 1. Additionally, noise sampled from a uniform and gaussian distributions are added to each time point to simulate real-world measurement variability.

It is important to note that not all theoretical combinations of criteria are valid. Specifically, a combination is considered invalid if the specified locations of change points exceed the given data length. For example, if the data length is 100, it is not feasible to have a change point at position 150, as the data cannot accommodate such a change point. Therefore, we excluded such invalid combinations from our analysis. This explains why some plots are missing in the visualization of the benchmarking results. In other words, the positions of change points must be within the range of the data length. Mathematically, if the maximum data length is *D*, then all change points must satisfy the condition *Change_Point_Position* ≤*D*. Figure 7 provides an example of such a dataset, generated with two change points (50, 100), 160 data points, and uniform noise. As shown in the figure, MICA successfully detects both change points.

**Fig. 7:**
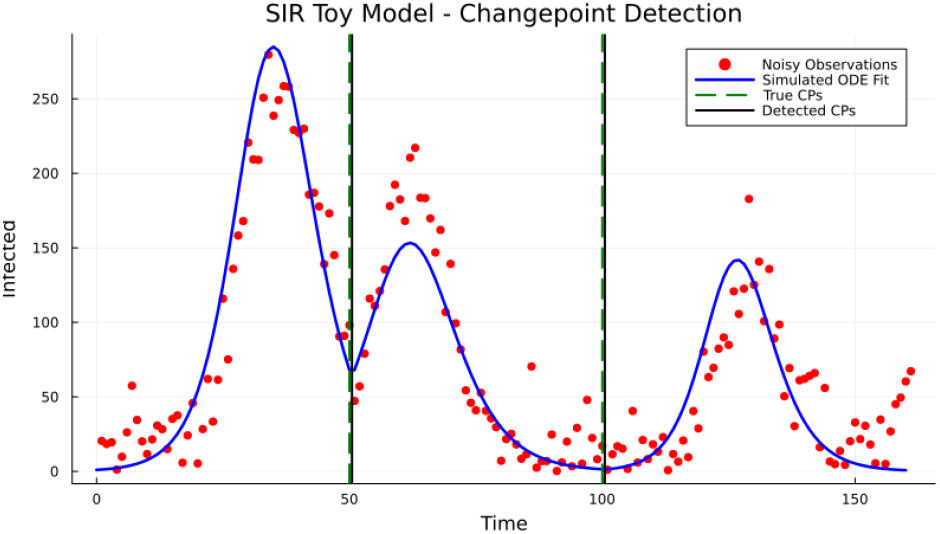
Application of MICA to noisy toy dataset generated by SIR model with two change points, data length 150

#### 3.1.2 Synthetic Benchmarking Results

Benchmarking performance is evaluated using a segment-wise loss function,

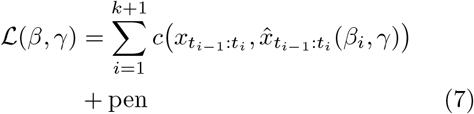

Where 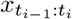 denotes the observed infected counts and 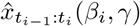 denotes the corresponding model simulations over segment *i* using Equations 6. Here, *β*_*i*_ is a segment-specific transmission parameter allowed to vary across segments, while *γ* is a non-segment-specific parameter shared across all segments. The function *c*(·,·) denotes the root mean square error (RMSE) computed within each segment.

The penalty term controls model complexity and is defined as

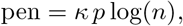

where *κ* is a penalty coefficient, *p* is the number of segment-specific model parameters, and *n* is the total number of data points. This formulation balances goodness of fit against model complexity, discouraging over-segmentation. The same loss and penalty structure is applied consistently across all synthetic experiments.

To quantify detection performance, we compute Recall, Precision, and the F1 Score, with an emphasis on Recall as the most relevant metric for identifying all true change points. Figures 8-10 illustrate accuracy trends under Gaussian and Uniform noise. MICA consistently achieved high recall across a broad range of noise levels and penalty configurations, particularly when penalty values were appropriately tuned. Additionally, the runtime associated with each combination of controlled experimental conditions is presented in Figure 11, illustrating the algorithm’s computational efficiency under varying scenarios. Across all tested configurations, typical runtimes ranged from approximately 40 to 200 seconds, depending on the number of change points, time points, and noise level.

**Fig. 8:**
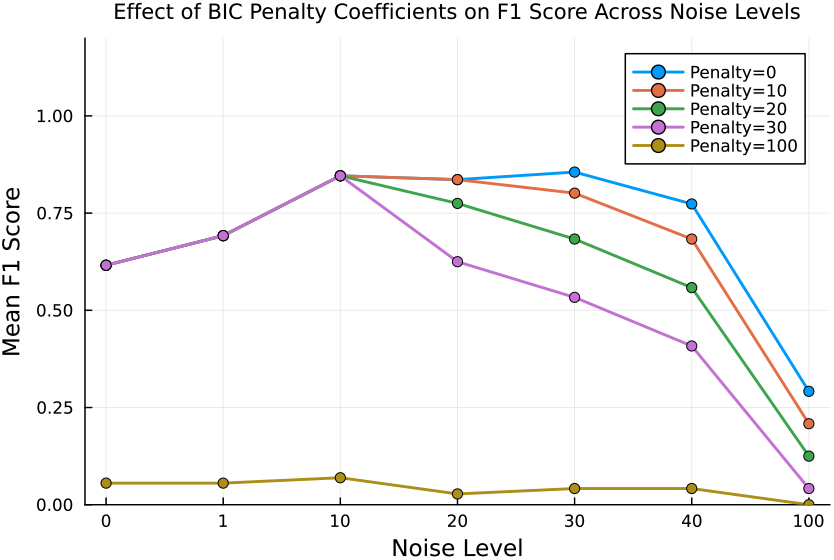
F1 Score vs. Noise Levels for Various Penalty Coefficients Using BIC-Based Penalty Scheme

**Fig. 9:**
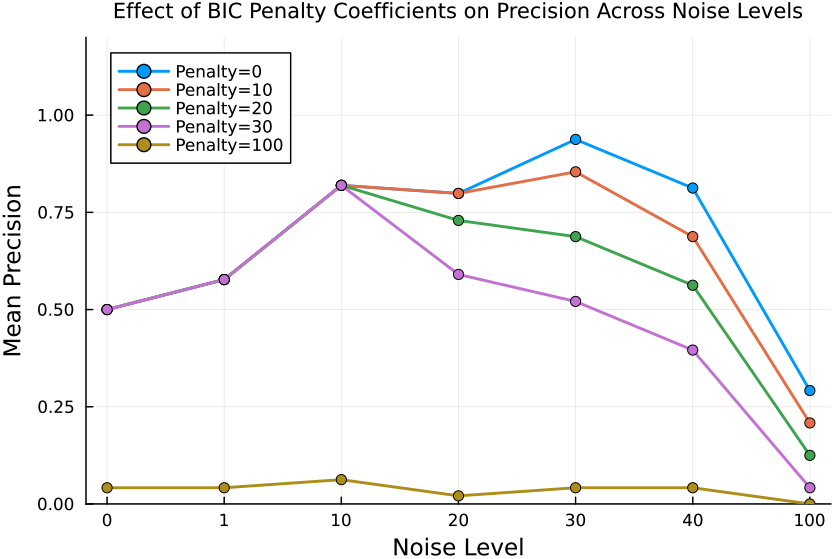
Precision vs. Noise Levels for Various Penalty Coefficients Using BIC-Based Penalty Scheme

**Fig. 10:**
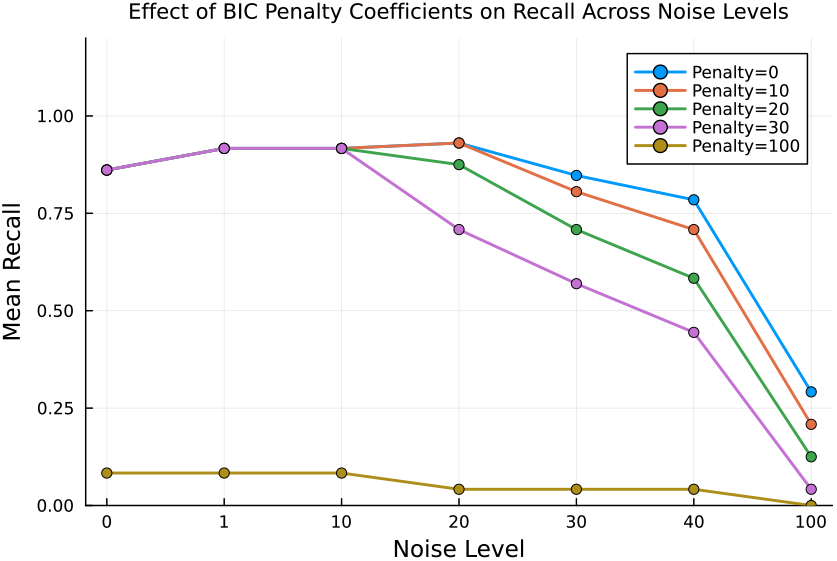
Recall vs. Noise Levels for Various Penalty Coefficients Using BIC-Based Penalty Scheme

**Fig. 11:**
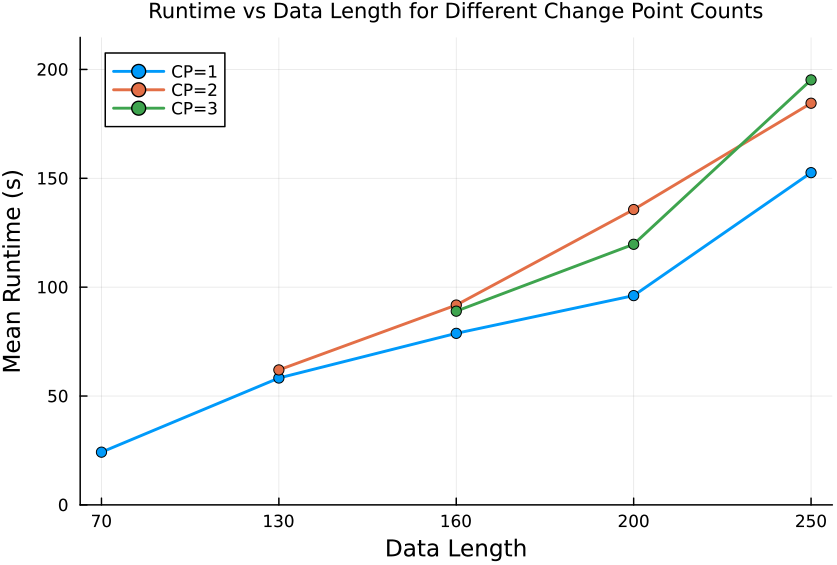
runtime vs data length for different numbers of change points

Regarding runtime, we recorded the number of numerical iterations performed by the ODE solver. Each simulation required approximately 3.7 × 10^4^ accepted steps and 2.2 × 10^5^ function evaluations, which represents the main numerical cost of our benchmarking. As the data length increases, more candidate segments must be evaluated, and thus more simulations are required. Consequently, the overall runtime is primarily driven by the number of solver iterations accumulated across all segments, rather than by changes in the cost of individual simulations. It is important to note that these benchmarks represent a worst-case scenario, since the chosen model does not admit an analytical solution and must be integrated numerically. For models with closed-form solutions, runtimes would be substantially lower, even though the scaling with respect to data length and change points would remain the same.

Overall, the synthetic evaluation confirms that MICA is capable of accurately detecting change points even in noisy or short time series, provided that the penalty term is properly calibrated.

### 3.2 MICA in COVID-19 Modeling

We applied MICA, to COVID-19 epidemiological data from Germany spanning January 27, 2020, to March 2, 2021, hence capturing the first two waves in Germany. We employed a system of differential equations incorporating 11 compartments, each representing distinct states in the disease progression. The variable *S* denotes susceptible individuals at risk of infection. *E*_0_ and *E*_1_ represent individuals in the early and secondary latent (exposed but not yet infectious) stages, respectively. The compartment *I*_0_ accounts for asymptomatic infected individuals, while *I*_1_ captures symptomatic cases. Individuals requiring regular hospitalization are represented by *I*_2_, and those admitted to intensive care units are categorized under *I*_3_. Recovered individuals, who are assumed to have acquired immunity, are represented by *R*, and deceased individuals are denoted by *D*. In addition to these, *C* represents the cumulative number of infections, and *V* denotes the cumulative number of vaccinated individuals. This compartmental structure provides a comprehensive framework for understanding the temporal dynamics of the pandemic and assessing the impact of interventions. Table 2 summarizes the model parameters, with the final column indicating whether each parameter is Segment-Specific (SP) or Non-Segment-Specific (NSP). Segment-Specific parameters are allowed to vary across detected change points, enabling the model to adapt to temporal shifts in disease dynamics, while Non-Segment-Specific parameters remain constant across all segments.

**Table 2:** Model parameters and their classification as segment-specific (SP) or non-segment-specific (NSP).

| Parameter | Explanation | SP/NSP |
| --- | --- | --- |
| $\beta$ | Transmission rate of the disease | SP |
| $\omega$ | Rate of loss of immunity | NSP |
| $\nu$ | Vaccination rate | SP |
| $\epsilon_0$ | Transition rate from $E_0$ to $E_1$ | NSP |
| $\epsilon_1$ | Transition rate from $E_1$ to $I_0$ | NSP |
| $\gamma_0$ | Recovery rate from asymptomatic infection ($I_0$) | NSP |
| $\gamma_1$ | Transition rate from symptomatic infection ($I_1$) to hospitalization or recovery | NSP |
| $\gamma_2$ | Transition rate from regular hospitalization ($I_2$) to ICU hospitalization ($I_3$) | NSP |
| $\gamma_3$ | Recovery rate from ICU hospitalization ($I_3$) | NSP |
| $p_1$ | Proportion of $E_1$ developing symptomatic infection ($I_1$) | SP |
| $p_{12}$ | Proportion of $I_1$ requiring regular hospitalization ($I_2$) | SP |
| $p_{23}$ | Proportion of $I_2$ requiring ICU care ($I_3$) | SP |
| $p_{1D}$ | Proportion of $I_1$ resulting in death | SP |
| $p_{2D}$ | Proportion of $I_2$ resulting in death | SP |
| $p_{3D}$ | Proportion of $I_3$ resulting in death | SP |

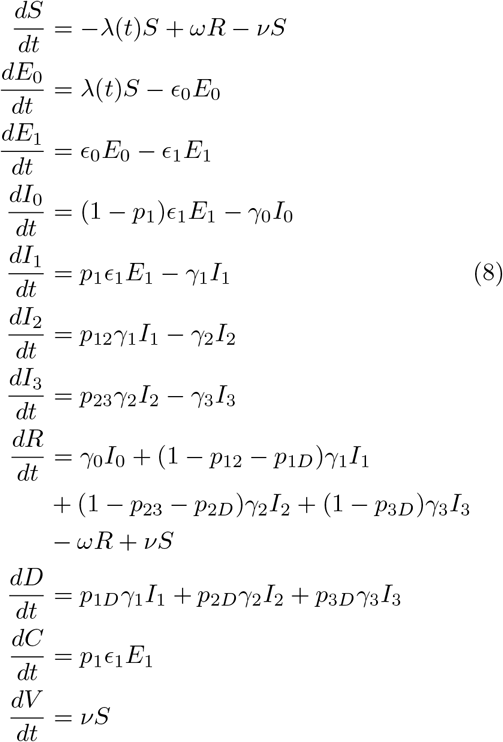

The force of infection is defined as

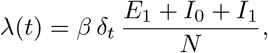

where *β* denotes the baseline transmission rate and *δ*_*t*_ is a seasonality factor given by

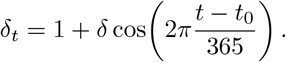

Here, *δ* represents the amplitude of seasonal variation, *t* denotes time (in days) and *t*_0_ is a reference time, set to 0 by default.

Furthermore, the total population *N* is given by:

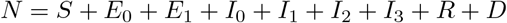

Change points in this context reflect structural transitions, such as the impact of public health interventions, shifts in population behavior, or the emergence of viral variants. MICA extends standard compartmental models by allowing selected parameters to vary across time segments, thereby capturing the evolving dynamics of the pandemic more effectively. For example, a particular lock-down measure in schools may change contact rates and, thus, infection rates between segments, but will not affect recovery rates of infected individuals, whereas e.g. a new viral variant will affect different sets of model parameters.

An objective function is designed to assess the goodness of fit between the observed real-world data and the simulated data generated by the model. The available data includes infected cases, hospitalizations, ICU admissions, deaths, and vaccination data. On the modeling side, we utilize Equations 8 to simulate the corresponding datasets. The objective function (ℒ) for fitting the model is therefore formulated as follows:

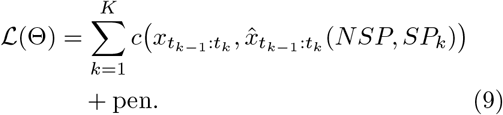

Here, 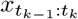 denotes the observed data and 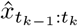 (*NSP, SP*_*k*_) denotes the corresponding model simulation obtained by solving Equations 8 using the non-segment-specific parameters *NSP*, which are shared across all segments, and the segment-specific parameter set *SP*_*k*_ associated with segment *k*, as summarized in Table 2. The function *c*(·) denotes the mean absolute error (MAE) computed within each segment.

To control the number of detected segments and discourage over-segmentation, a Bayesian Information Criterion (BIC)–inspired penalty was employed. The penalty term is defined as

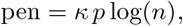

where *p* denotes the number of segment-specific parameters and *n* is the total number of data points. The constant *κ* = 40.0 was selected empirically by evaluating a small range of values and choosing the one that produced stable and consistent segmentation results across repeated analyses. This choice was based on empirical performance rather than formal optimization.

Figure 12 presents the results of change point detection obtained using MICA. The top panel compares observed and simulated data for reported infections, hospitalizations, ICU admissions, and deaths. Detected change points are marked by vertical green lines. The bottom panel shows the relative change to the first segment (RCFS) for each segment-specific parameter: detection rate (*p*_1_), infection rate (*β*), hospitalization rate (*p*_12_), ICU admission rate (*p*_23_), and stage-specific fatality rates (*p*_1*D*_, *p*_2*D*_, *p*_3*D*_).

**Fig. 12:**
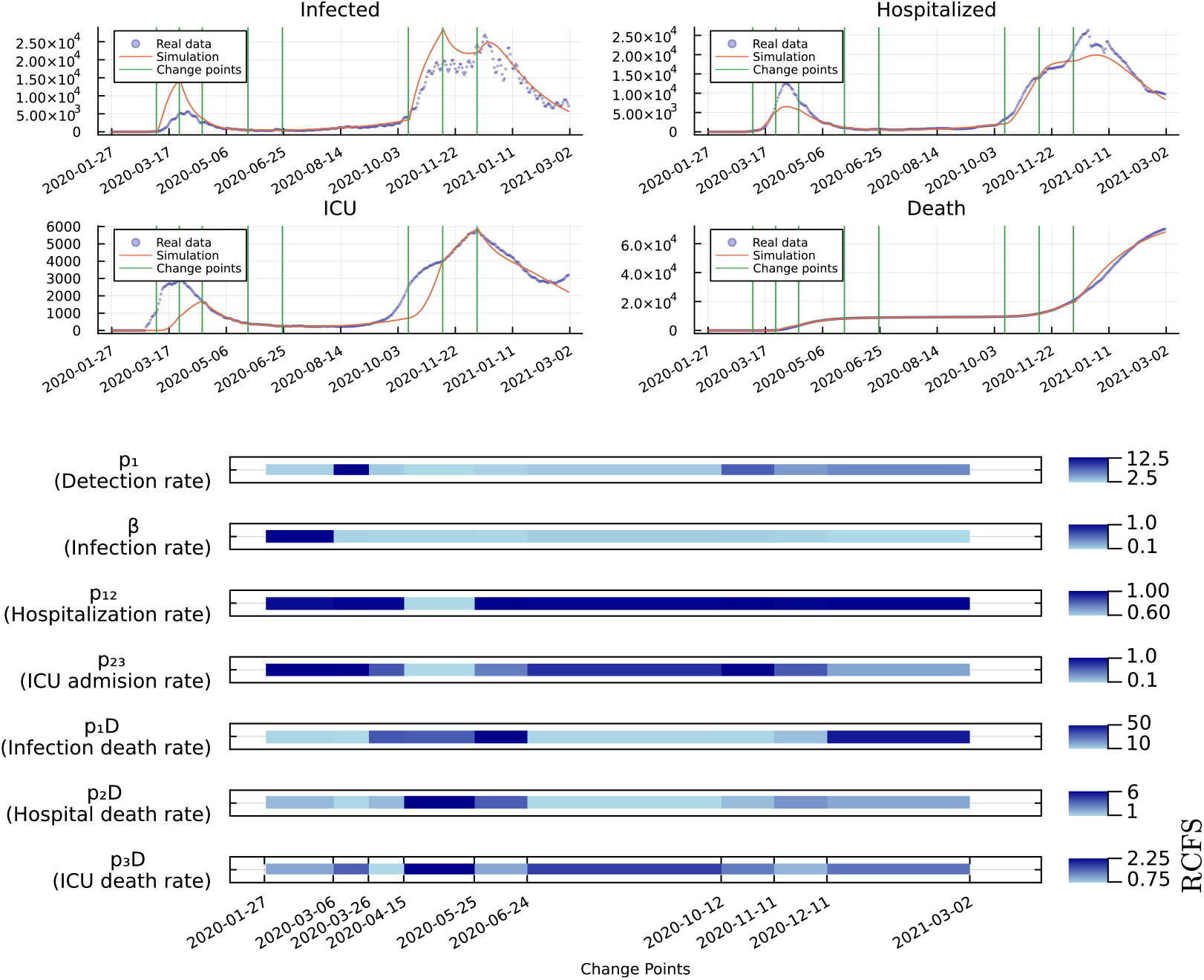
Simulated COVID-19 dynamics in Germany with detected change points and segment-specific parameter shifts. Top: Observed vs. simulated infections, hospitalizations, ICU admissions, and deaths. Vertical green lines indicate detected change points. Bottom: Relative change of segment-specific parameters with respect to the first segment (RCFS), revealing shifts in transmission, hospitalization, and fatality dynamics over time.

Eight change points were detected in approximately 90 minutes, dividing the epidemic timeline into nine distinct regimes. These regimes correspond to periods marked by major policy interventions, behavioral adjustments, and epidemiological shifts, with each transition reflected in parameter dynamics. The first change point, on March 6, 2020, coincided with border closures and the cancellation of large gatherings. The second, on March 26, 2020, followed nationwide lockdown measures including school closures, closure of churches, cancellation of sports events, and restrictions on non-essential shops. The third change point, on April 15, 2020, aligned with the introduction of stricter distancing measures and the Corona-Warn-App. The fourth, on May 25, 2020, corresponded to the reopening of schools and sports activities. The fifth, on June 24, 2020, coincided with broader relaxation of distancing rules and border reopening. The sixth, on October 12, 2020, was observed just prior to stay-at-home orders. The seventh, on November 11, 2020, matched the implementation of mask mandates in schools and partial lockdowns. Finally, the eighth change point, on December 11, 2020, occurred immediately before renewed school closures and full lockdowns.

Overall, the main parameters followed expected patterns across these phases. Transmission (*β*) decreased sharply after the first interventions in March, falling to less than 5% of baseline after border closures and further after nationwide lockdowns. Subsequent measures in April sustained low transmission, while partial and broader relaxations from late May and June were accompanied by stable or slightly higher *β*, followed by a clear rebound during the summer. The renewed restrictions in autumn again suppressed transmission, and the December lockdown brought *β* to its lowest recorded level (around 2% of baseline). Detection (*p*_1_) also tracked intervention phases. It rose strongly after the early March measures, reflecting rapid expansion of testing, dipped somewhat after the March 26 lockdown, and declined further into April. With reopening from May onward, detection recovered, reached higher levels during the summer, and peaked in October at over eight times the baseline. Detection remained elevated through the subsequent restrictions in November and December. Hospitalization (*p*_12_) and ICU admission (*p*_23_) probabilities showed more modest variation but were broadly consistent with intervention timing. These parameters were stable during the first lockdown, declined in April, and then increased with the partial and broader reopening phases. After October, hospitalization returned to baseline values, while ICU admission fell again under the later restrictions. Fatality parameters (*p*_1*D*_, *p*_2*D*_, *p*_3*D*_) showed more variability across phases. In general, mortality ratios decreased relative to baseline after the first interventions, rose during certain later phases, and remained elevated in December despite strong suppression of transmission.

While these broad trends align with expectations, not all parameter shifts followed the anticipated pattern. For example, infection and hospital fatality occasionally increased even during phases when transmission was falling or detection was rising, and ICU fatality sometimes moved in the opposite direction from the other mortality measures. Beyond the model’s internal mechanics, these patterns might also reflect real-world processes such as reporting delays, changes in who was being tested, or variation in the severity of illness over time.

### 3.3 MICA in Wind Turbine Monitoring

Wind turbines operate under variable environmental and load conditions, making early detection of internal behavioral shifts essential for reliability and predictive maintenance. While status logs record events like start-ups and shutdowns, they offer limited insight into how such conditions impact internal dynamics, particularly generator cooling behavior influenced by wind speed, ambient temperature, and thermal resistance. In this study, we apply MICA to thermal model parameters to identify abrupt changes in cooling dynamics that may indicate operational faults or emerging component degradation. By aligning detected change points with turbine status events, we aim to enhance fault diagnostics and enable more informed maintenance strategies.

Several recent studies have explored fault detection in wind turbines using Supervisory Control and Data Acquisition (SCADA) data, employing approaches such as thermophysics-based diagnostics and kernel-based statistical methods [20–22]. While these methods provide valuable insights, they often lack adaptability in model structure, particularly in supporting dynamic parameter updates or accommodating global parameter consistency alongside local variability. Many rely heavily on the statistical properties of the data without integrating mechanisms for dynamic model refinement. In contrast, MICA addresses this gap by enabling selective parameter adaptation across temporal segments: certain parameters are held constant to maintain global stability, while others are allowed to vary to capture local behavioral changes. This hybrid structure improves both modeling flexibility and fault detection accuracy in operational settings.

The Kelmarsh Wind Farm SCADA dataset was used as the second application of MICA. The dataset was released by Cubico Sustainable Investments Ltd. under a CC-BY-4.0 license and exported via the Greenbyte platform on January 27, 2022 [14]. The dataset comprises 10-minute resolution SCADA and event log data from six Senvion MM92 turbines in the UK, covering the period from 2016 to mid-2021. For this study, we focused on a single turbine, Kelmarsh 1, over an 18-day window from January 1 to January 18, 2021. The analysis used three primary signals, generator temperature, wind speed, and ambient temperature, all recorded at 10-minute intervals. These were used as inputs to the thermal model and change point detection framework.

The mathematical model employed in this application is based on the “Model Sensor” framework by Zhang et al. [19], which captures the dynamic relationship between generator temperature and environmental inputs. The key idea is to estimate generator temperature based on energy balance principles, accounting for copper losses and heat dissipation via thermal resistance. Figure 13 illustrates this conceptual model.

**Fig. 13:**
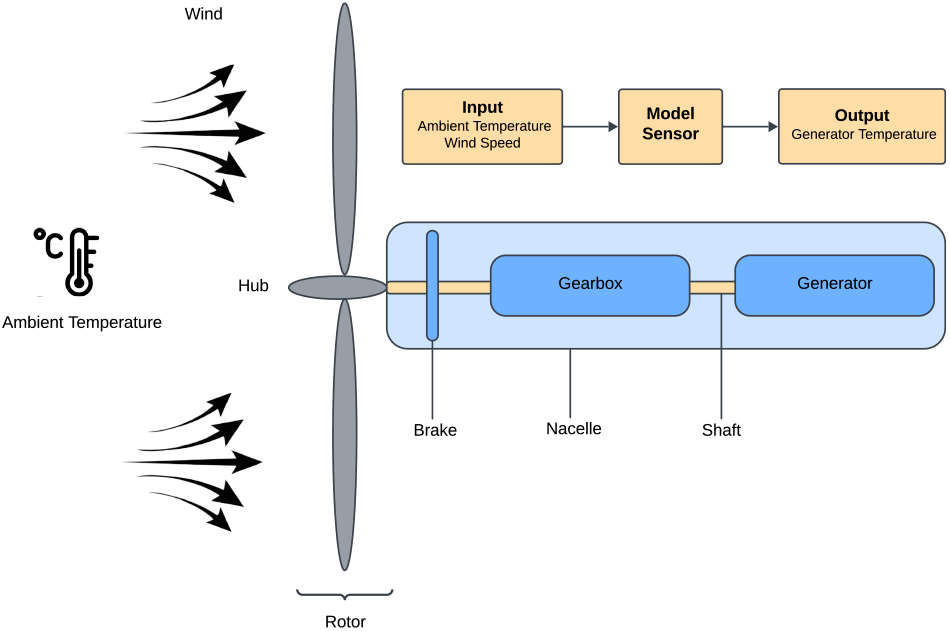
This figure depicts the key components of a wind turbine, including the blades, gearbox, yaw system, and generator winding. The focus is on the relationship between the generator temperature (model output) and two model inputs: ambient temperature and wind speed. The ”Model Sensor” illustrates the dynamic relationship between these inputs and the output, highlighting how changes in the ambient conditions and wind speed affect the generator’s temperature.

The model captures the relationship between the generator temperature *T*_*g*_, wind speed *V*_*w*_, and ambient temperature *T*_*a*_. This relationship is governed by the energy balance equation [16], which considers the energy input *Q*_in_ due to copper losses and the energy output *Q*_out_ due to cooling:

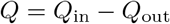

The generator winding temperature change (Δ*T*) is related to the energy *Q* and thermal capacitance *C*_*g*_ as [15]:

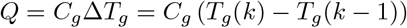

where *T*_*g*_(*k*) represents the generator winding temperature at time *k*.

It is considered that the nonlinear relationship between copper losses and wind speed *V*_*w*_, which can be approximated by a third-degree polynomial [17]:

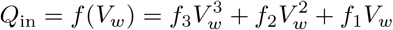

Similarly, the heat conduction due to cooling, *Q*_out_, is modeled considering the thermal resistance *R*_*ga*_ [15], which depends on wind speed through another third-degree polynomial [18]:

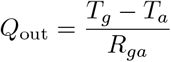

where

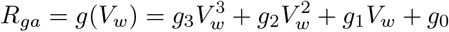

The generator temperature is thus given by:

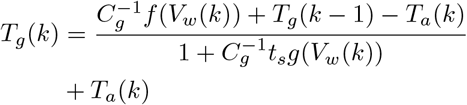

By substituting *f* (*V*_*w*_(*k*)) and *g*(*V*_*w*_(*k*)) from previous equations, we obtain the final model that describes the relationship between generator temperature, ambient temperature, and wind speed. This model is expressed as:

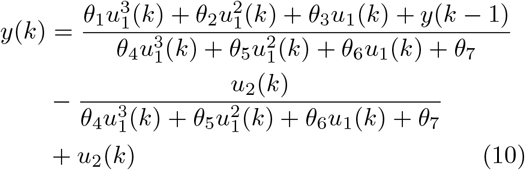

Where 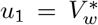 and 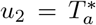 are the model inputs, and 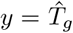 is the model output. And the parameter vector ***θ*** is defined as:

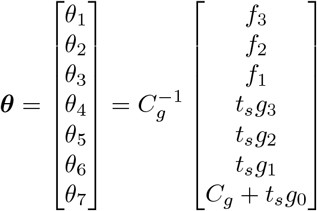

where ***θ*** is the vector of model sensor parameters that need to be updated regularly for wind turbine change point detection.

We classify model parameters into segment-specific (SP) and non-segment-specific (NSP) types, as detailed in Table 3. Parameters related to copper losses, specifically those governing the relationship between wind speed and electrical heating, are assumed to be constant across all segments. This reflects the assumption that electrical losses due to current flow in the windings remain relatively stable regardless of operational changes or fault conditions. In contrast, parameters associated with thermal resistance are treated as segment-specific, allowing the model to adapt to varying cooling dynamics influenced by operational or environmental changes. For example, cooling efficiency may vary with fluctuations in wind speed, ambient temperature, or turbine state (e.g., during startup or fault recovery). Allowing thermal resistance parameters to vary across segments enhances the model’s ability to capture these dynamic effects and improves the precision of change point detection. The full classification of parameters is summarized in Table 3, where SP denotes segment-specific parameters and NSP denotes those held constant across the entire time series.

**Table 3:** Model parameters and classification into segment-specific (SP) and non-segment-specific (NSP)

| Parameter Name | Parameter Description | Type |
| --- | --- | --- |
| $\theta_1$ | Coefficient related to copper loss polynomial | NSP |
| $\theta_2$ | Coefficient related to copper loss polynomial | NSP |
| $\theta_3$ | Coefficient related to copper loss polynomial | NSP |
| $\theta_4$ | Coefficient related to thermal resistance polynomial | SP |
| $\theta_5$ | Coefficient related to thermal resistance polynomial | SP |
| $\theta_6$ | Coefficient related to thermal resistance polynomial | SP |
| $\theta_7$ | Coefficient related to thermal resistance polynomial | SP |

To estimate the parameters of the model in Equation 10, we define the following objective function:

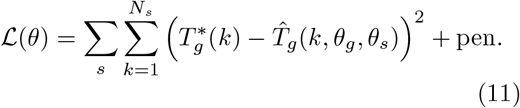

Here, 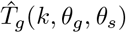 denotes the model-predicted generator temperature obtained by solving Equation 10 at time point k, where *θ*_*g*_ represents the non-segment-specific parameters and *θ*_*s*_ denotes the segment-specific parameters associated with segment *s*, as detailed in Table 3. The quantity 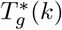 denotes the observed generator temperature from the SCADA dataset at time point k, and *N*_*s*_ is the number of observations within segment *s*.

The penalty term in this application followed a BIC-based formulation to regulate the number of detected segments and to mitigate over-segmentation. Specifically, we used

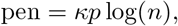

where *p* is the number of segment-specific parameters and *n* denotes the data length. The constant *κ* = 100.0 was determined empirically after testing several values and selecting the one that produced stable and interpretable segmentations across repeated runs. This choice was made through empirical assessment rather than formal optimization.

Figure 14 presents the results of thermal modeling and change point detection using MICA. The top subplot shows detected change points superimposed on the actual and simulated generator temperatures. The bottom subplot illustrates the relative change in segment-specific parameters (*θ*_1_–*θ*_4_) compared to the first segment. These parameters represent thermal resistance coefficients and reflect changes in the cooling dynamics of the generator.

**Fig. 14:**
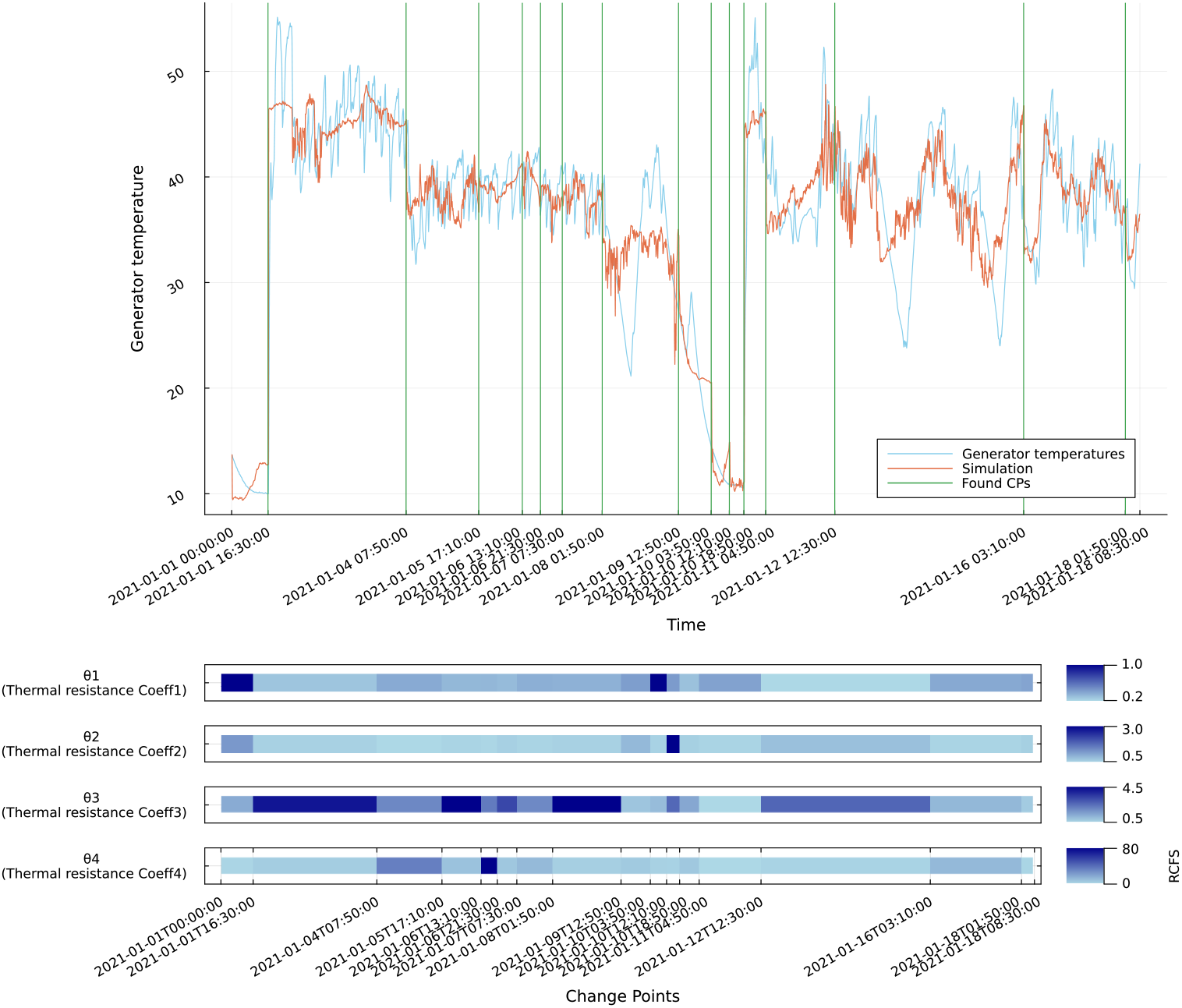
Detected change points and their effect on generator thermal model parameters. Top: actual and simulated temperatures with change points. Bottom: relative shifts in segment-specific thermal parameters across segments.

Fifteen change points were detected within approximately 34 minutes of computation for a data length of 2,500 time points. The first change point (2021-01-01 16:30) coincides with a startup sequence involving mains run-up and gearbox warm-up. This is marked by a sharp drop in *θ*_1_ and *θ*_2_ and a simultaneous increase in *θ*_3_ and *θ*_4_, indicating a transition from stationary to active thermal behavior. Several subsequent change points (e.g., 2021-01-04 07:50, 2021-01-05 17:10) occur in the absence of status log entries but feature spikes in *θ*_4_, suggesting undocumented or latent thermal shifts.

Other change points align with known events such as external stops due to low wind (e.g., 2021-01-07 07:30, 2021-01-08 01:50) and icing conditions (e.g., 2021-01-09 12:50). These typically show transient increases in *θ*_2_ and *θ*_3_ and varying responses in *θ*_4_, reflecting changes in heat dissipation and cooling response.

The final segments exhibit more stable behavior, with *θ* parameters returning closer to baseline or oscillating within narrower ranges. However, certain transitions (e.g., 2021-01-12 12:30) show extreme drops in multiple parameters, possibly due to short-term signal artifacts or local data irregularities.

Overall, the alignment of parameter shifts with turbine events supports the model’s capacity to detect and explain operational transitions. The relative parameter changes provide interpretable physical meaning, indicating when and how the generator’s thermal response is altered by external conditions or internal faults.

The detected change points were systematically compared with turbine status logs contained in the SCADA dataset to assess their alignment with known operational events. As shown in Table 4, several change points closely aligned with startup sequences, including run-up, mains connection, and gearbox warm-up stages, indicating a strong correspondence between physical thermal transitions and externally reported events. Others occurred shortly before external stops, particularly due to low wind conditions, suggesting early indicators of shutdown behavior. A subset of change points exhibited no clear status association, highlighting either undocumented anomalies or latent physical transitions not captured by the status system. This analysis supports the hypothesis that change point detection applied to thermal model parameters can effectively identify both recorded and unrecorded shifts in operational behavior.

**Table 4:** Alignment between detected change points and turbine status events.

| Change Point | Nearest Status Event | Alignment | Event Type | Description |
| --- | --- | --- | --- | --- |
| 2021-01-01 16:30 | 2021-01-01 16:46 | Strong | Startup sequence | Change point closely precedes a full startup sequence, indicating a thermal transition as the turbine returns to operation. |
| 2021-01-04 07:50 | 2021-01-03 22:20 | Weak | Data communication unavailable | No direct operational event; data communication issue recorded 9 hours earlier. |
| 2021-01-05 17:10 | 2021-01-06 23:04 | Weak | External stop and restart | Change point occurs more than 24 hours before an external stop/start cycle; not closely aligned. |
| 2021-01-06 13:10 | 2021-01-06 23:04 | Weak | External stop and restart | Occurs 10 hours before a stop/start cycle. Weak alignment. |
| 2021-01-06 21:30 | 2021-01-06 23:04 | Moderate | External stop and restart | Occurs 1.5 hours before turbine stop. Possible lead-in to shutdown. |
| 2021-01-07 07:30 | 2021-01-07 01:43 | Moderate | Frequent external stops | Change point occurs during a dense period of operational cycling; moderate alignment with thermal dynamics. |
| 2021-01-08 01:50 | 2021-01-08 03:27 | Moderate | External stop (low wind) | Change point occurs 1.5 hours before shutdown. Potential early cooling anomaly. |
| 2021-01-09 12:50 | 2021-01-09 16:26 | Weak | Run-up and cycling | Occurs 3.5 hours before operational activity. Not directly aligned. |
| 2021-01-10 03:50 | 2021-01-10 07:54 | Weak | Reset of icing condition | Occurs 4 hours prior to icing status reset. Unclear linkage. |
| 2021-01-10 12:10 | 2021-01-10 18:48 | Weak | Startup after icing | Occurs 6.5 hours before restart. Weak alignment. |
| 2021-01-10 18:50 | 2021-01-10 18:50 | Strong | Gearbox warm-up | Change point aligns exactly with turbine restart and gearbox warm-up sequence. |
| 2021-01-11 04:50 | 2021-01-10 18:50 | Moderate | Extended operation | Follows a long stable run. Possible drift or thermal regime shift. |
| 2021-01-12 12:30 | 2021-01-12 15:42 | Moderate | External stop (low wind) | Change point precedes external stop and restart cycle by 3 hours. |
| 2021-01-16 03:10 | 2021-01-15 15:30 | Weak | Multiple stops | Change point occurs during period of many external stops; no clear triggering event. |
| 2021-01-18 01:50 | 2021-01-19 12:24 | Weak | Yaw and battery test | Occurs more than a day before status activity. No strong alignment. |

## 4 Discussion

The results presented in this study demonstrate the effectiveness and versatility of MICA, our proposed change point detection algorithm, across a variety of domains. By enabling selective parameter adaptation, MICA allows dynamic models to incorporate structural changes in a principled and interpretable manner. Applied to both synthetic datasets and real-world systems, including epidemiological modeling of COVID-19 and fault detection in wind turbine cooling systems, MICA successfully identified significant temporal shifts aligned with known events or operational anomalies.

A central strength of MICA lies in its ability to distinguish between segment-specific and non-segment-specific parameters. This feature offers a major advantage over conventional approaches that assume either full parameter constancy or complete variability. In the COVID-19 modeling scenario, this allowed the model to adapt to shifts in disease transmission while preserving invariant biological parameters. In the wind turbine application, it enabled detection of cooling efficiency changes without incorrectly attributing noise to stable electrical losses.

Compared to existing change point detection methods, such as kernel-based techniques, fixed-threshold statistical models, or manually defined intervention points, MICA introduces a flexible and model-aware strategy. It integrates seamlessly with dynamic systems and enhances both the accuracy and explanatory power of simulation results.

Nevertheless, the method has certain limitations. The selection of the penalty term significantly influences the granularity of detected change points; inappropriate tuning can lead to either over-segmentation or missed transitions.. Moreover, alignment with real-world event logs (such as SCADA status files) may be imperfect due to sensor delays, missing data, or asynchronous reporting. Additionally, the current use of a binary segmentation introduces a greedy search heuristic, which, while efficient, may not guarantee globally optimal segmentation. Future work will focus on improving robustness under uncertainty, supporting a broader class of dynamic models, and incorporating more general, non-greedy segmentation strategies to enhance detection accuracy and flexibility.

Despite these challenges, MICA shows strong potential for generalization. Its domain-agnostic formulation makes it applicable to a wide range of time-dependent systems, including industrial monitoring, biomedical signal processing, environmental sensing, and econometrics. Future directions include the development of automated penalty tuning methods, real-time streaming implementations, and integration with more complex or multivariate system models.

Overall, MICA provides a flexible, interpretable, and computationally viable framework for structural change detection in dynamic systems. Its capacity to operate across domains while respecting the inherent structure of underlying models positions it as a valuable tool for both scientific investigation and practical decision-making.

## 5 Conclusion

We presented MICA, a flexible and model-driven algorithm for detecting change points in time series governed by dynamic systems. Unlike traditional statistical approaches, MICA operates directly on the model’s structure by allowing selective parameter segmentation, capturing changes in system dynamics while preserving parameter stability where appropriate.

By combining a tailored binary segmentation strategy with an adaptive optimization procedure, MICA enables precise and interpretable identification of structural shifts in both synthetic and real-world datasets. Its application to COVID-19 epidemiological modeling and wind turbine operational monitoring illustrates its effectiveness in revealing meaningful change points aligned with known interventions and operational events.

The algorithm’s generality, demonstrated across domains with distinct system characteristics, highlights its utility for a wide range of applications, from public health modeling to industrial fault detection. As a domain-agnostic tool with built-in model awareness, MICA contributes a novel, versatile approach to structural change analysis in dynamic systems.

## 6 Acknowledgments

We thank the members of the Kaderali group, in particular Dr. Rahul Brahma and Yasas Wijesekara, for their valuable comments and suggestions during the preparation of this study. We also thank Johannes Apelt for his input on the development of the method package in Julia. Their feedback greatly contributed to improving the quality of this work.

## Declarations

### Author Contributions

ML developed the methodology, implemented the computational framework, carried out the analyses, and drafted the manuscript. LK conceived the study, contributed to the design of the methodology, guided data interpretation, and contributed to manuscript preparation. Both authors reviewed, edited, and approved the final version of the manuscript.

### Data Availability

The wind turbine modeling data used in this study are publicly available via Zenodo at https://doi.org/10.5281/zenodo.5841834. The COVID-19 epidemiological datasets were obtained from publicly accessible repositories maintained by the Robert Koch Institute (RKI), Germany, including COVID-19 mortality data (https://github.com/robert-koch-institut/COVID-19-Todesfaelle in Deutschland),seven-day incidence data (https://github.com/robert-koch-institut/COVID-19_Tage-Inzidenz_in_Deutschland), hospitalization data (https://github.com/robert-koch-institut/COVID-19-Hospitalisierungen_in_Deutschland), ICU capacity and occupancy data (https://github.com/robert-koch-institut/Intensivkapazitaeten_und_COVID-19-Intensivbettenbelegung_in_Deutschland), and vaccination data (https://github.com/robert-koch-institut/COVID-19-Impfungen_in_Deutschland).

The full source code and implementation used in this study are openly available at https://github.com/Mehdilotfi7/MICA.

